# Integrated analysis of multimodal long-read epigenetic assays

**DOI:** 10.1101/2025.11.09.687458

**Authors:** Jeremy Marcus, Oberon Dixon-Luinenburg, Nathan Gamarra, Jacob P. Schwartz, Michal Rozenwald, Annie Maslan, Fyodor D. Urnov, Aaron F. Straight, Nicolas Altemose, Nilah M. Ioannidis, Aaron Streets

## Abstract

Long-read sequencing assays that detect base modifications are becoming increasingly important research tools for the study of epigenetic regulation, especially with the development of DiMeLo-seq and similar methods that deposit non-native base modifications to mark a range of epigenetic features such as protein-DNA interactions and chromatin accessibility. A main benefit of these methods is their inherent capacity for multimodality, enabling the encoding of multiple genomic signals onto single nucleic acid molecules. However, there are limited tools available for visualization and statistical analysis of this type of multimodal data. Here we introduce *dimelo-toolkit*, a python package built to enable flexible visualizations and easy integration into custom data processing workflows. We demonstrate the utility of *dimelo-toolkit*’s preset visualizations of multiple base modifications in long-read single-molecule sequencing data with a novel extension of the DiMeLo-seq protocol that can capture three separate aspects of chromatin state on the same single reads: target protein binding, CpG methylation, and chromatin accessibility. We apply this multimodal method to simultaneously map chromatin accessibility, CpG methylation, and LMNB1 and CTCF binding patterns, respectively, in GM12878 cells. Our flexible design allows us to investigate previously unexplored technical biases that arise when working with this type of multimodal data. Additionally, we show that *dimelo-toolkit* enables analysis for a wide range of other long-read sequencing methods, such as mapping endogenous patterns in RNA base modifications with direct RNA sequencing. This software tool will pave the way for developing well-optimized protocols and help unlock previously inaccessible biological insights.

## Introduction

The genome in higher eukaryotes encodes a vast set of distinct cellular phenotypes, requiring sophisticated machinery to regulate gene activity and maintain cell state. This epigenetic regulation relies on the coordination of direct nucleotide modifications in the form of DNA methylation, binding and modification of proteins such as transcription factors and histones, and larger-scale structural features including chromatin accessibility and topologically associated domains^1^. Genome-wide measurement of these epigenetic features has proven transformative for understanding the structure and function of the genome^2–4^. However, performing multiple epigenetic measurements simultaneously in the same cells remains difficult using standard technologies.

The most widely used methods for assessing epigenetic state currently rely on short-read sequencing. Methods for measuring protein-DNA interactions or chromatin accessibility rely on fragmentation of the genome and subsequent fragment enrichment. Following amplification, sequencing, and alignment, sequencing coverage at genomic loci is used as a proxy for the phenomenon of interest. Protein-DNA interaction assays such as DamID^5,6^, ChIP-seq^7^, and CUT&TAG^8^ enrich for reads which are bound by a target protein of interest, while chromatin accessibility assays such as DNAse-seq^9^ and ATAC-seq^10^ enrich for reads from accessible regions of the genome. These short-read assays all rely on the same final output: the number of reads which map to each genomic location. As a result, they are mutually incompatible and must be performed separately. Additionally, short-read sequencing technologies have important limitations. These methods do not directly measure DNA modifications, instead requiring base conversion protocols such as bisulfite sequencing^11^ or EM-seq^12^. Furthermore, the short length of the sequenced fragments makes it difficult to map reads to repetitive sequences or perform high-confidence haplotype phasing.

A new class of assays utilize third-generation long-read sequencing and DNA modifications to interrogate epigenetic state. Assays such as DiMeLo-seq^13,14^, Fiber-seq^15^, SAMOSA^16^, nanoNOME-seq^17^, and others^18–27^ write information about protein-DNA interactions or chromatin accessibility directly onto the DNA itself through the introduction of base modifications in exogenous motif contexts proximal to phenomena of interest. These modifications can be detected directly alongside primary DNA sequence and endogenous methylation on reads many kilobases in length using Oxford Nanopore Technologies (ONT) or Pacific Biosciences HiFi sequencing technologies. Long-read data enables applications including quantitative haplotype-specific chromatin actuation, mechanistic understanding of the ultra-repetitive regions of the centromere, and assessment of coordinated binding events on single DNA molecules.

As these experimental methods have advanced, software tools to visualize and analyze this new data modality remain an area of active work. Visualizing local and global patterns and performing genome-wide statistical analyses requires tools beyond those typically used for read-coverage-based short-read assays. Furthermore, the wide array of possible exogenous modifications, such as methylation and hydroxymethylation, in different motif contexts, such as CpG, GpC, and CpC, requires a flexible and modular approach. We previously developed a software package for visualization of DiMeLo-seq data named *dimelo*^14^, which provided plotting tools hardcoded for adenine and CpG methylation. To support further development of modification-based long-read assays, we have expanded the scope of past software built for DiMeLo-seq and released the new *dimelo-toolkit*. By using *modkit*^28^ to parse sequencing data and then optimizing data loading and processing within our core package, key operations are sped up from hours or days with the previous tool to seconds or minutes with *dimelo-toolkit*. We have maintained the easy-to-use interfaces of the previous tool while implementing a new modular and flexible design to facilitate both end-to-end pipelines for high-quality visualizations and vectorized data extraction for more advanced workflows. Speeding up parsing and loading enables quick iteration in analysis, while the flexibility enables compatibility with a wide range of assays and pipeline configurations. We have also added the capability to process any type of base modification in any sequence context, including modifications found in RNA^29^ rather than DNA. Thus *dimelo-toolkit* has the potential to help researchers working with a wide range of different experimental methods and biological questions.

With these features, *dimelo-toolkit* facilitates the analysis of datasets containing an arbitrary number of different nucleotide modifications, creating the opportunity to build experimental methods which encode multiple epigenetic features onto single chromatin fibers. These multimodal assays enable study of the biological interactions between epigenetic phenomena, such as the function of co-occurring protein-DNA binding events or relationships between binding events and chromatin structure. Here, we leverage the multimodal capabilities of *dimelo-toolkit* to drive analysis for a new protocol that simultaneously measures protein-DNA interaction, chromatin accessibility, and CpG methylation as part of a single assay. We characterize these multiplexed measurement signals and correct for signal crosstalk, thus enabling high-read-depth, single-molecule, and haplotype-resolvable information for all three signals.

## Results

### *dimelo-toolkit* workflow

*dimelo-toolkit* defines a systematic workflow for working with modified nucleotide data in python and producing publication-quality visualizations (Figure 1). This allows users to easily interpret and analyze the results of experiments which rely on nucleotide modification state as the primary output modality, whether through measurement of endogenous modification patterns or the introduction of exogenous modification signals.

**Figure 1.**
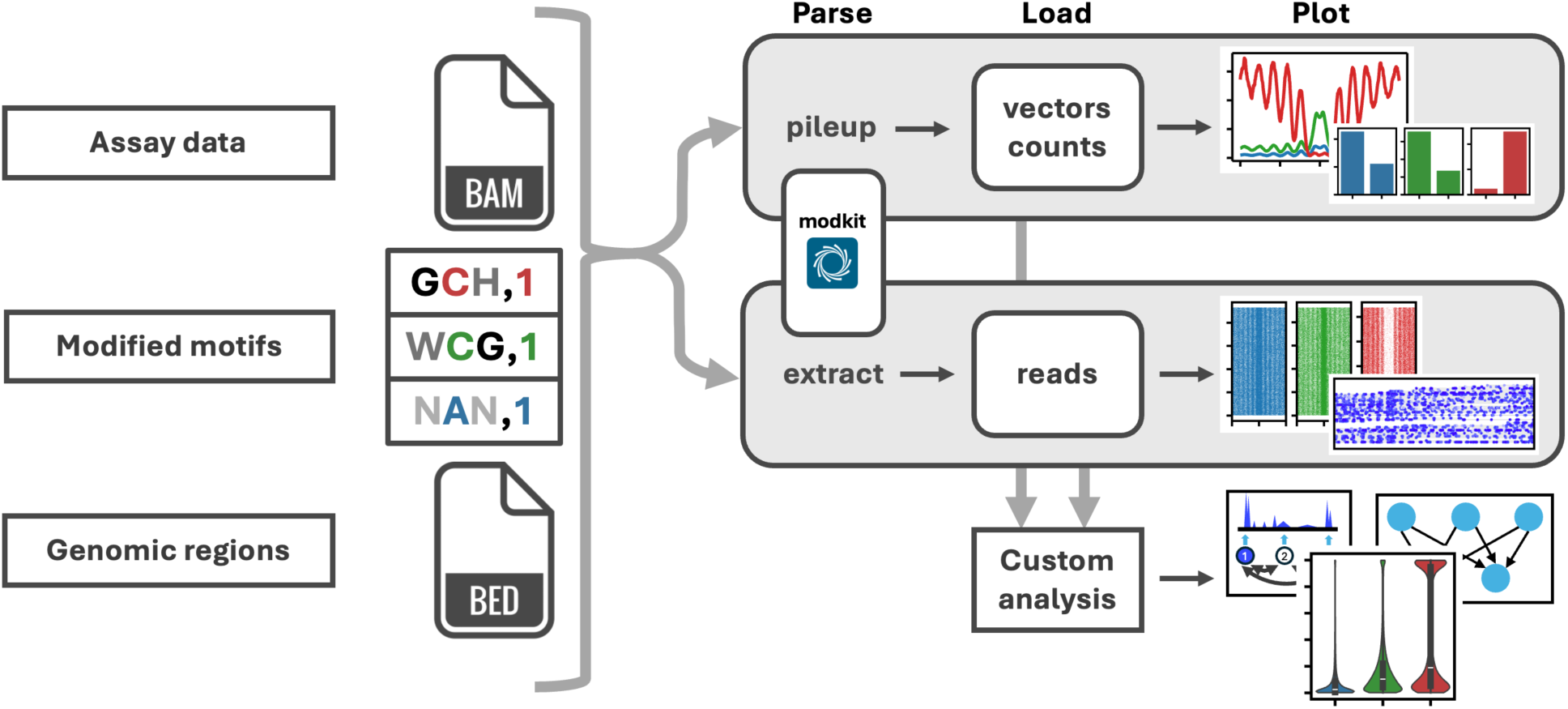
dimelo-toolkit workflow. dimelo-toolkit accepts base-called, modification-tagged, and aligned sequencing data in a BAM format file as primary input, along with base modification codes and sequence contexts for an arbitrary number of modification types. The user may specify genomic regions of interest via a BED file or region strings. Given these inputs, dimelo-toolkit can either collapse modification information into a single value for each genomic coordinate (pileup), representing the fraction of the reads covering each position which exhibit a given modification, or tabulate information for each read in each region (extract). Parsing operations create compressed and indexed intermediate files from outputs generated by ONT modkit, enabling quick iteration when prototyping new downstream analyses. The package provides methods for producing preset plots from these intermediate files. Additionally, the methods for loading parsed data are made available to the user to enable the building of custom python analysis pipelines, ranging from custom data visualizations to training models on large volumes of data.

As a primary input, the package accepts aligned and base-called long-read sequencing data in a BAM format file containing the appropriate base modification tags (MM and ML) for an arbitrary number of endogenous and exogenous modification types, such as 5mC and 6mA. To disambiguate between modification signals in different contexts (i.e, 5mC in CpG vs. GpC motifs, or 6mA specific to GATC motifs), the user can specify the genomic motifs at which to extract modification events using IUPAC standard nucleotide codes, a zero-indexed modification position, and optionally a mod code. For example, GCH,1,m specifies a motif with guanine at position 0, a 5-methylcytosine at position 1, and an adenine, cytosine, or thymine at position 2. Additionally, many of the methods in *dimelo-toolkit* allow the user to separately specify genomic regions of interest. Given these inputs, *dimelo-toolkit* defines two major workflow branches. In the pileup operation, reads are collapsed into a single value for each genomic coordinate, representing the fraction of the reads covering each position which exhibit a given modification. This enables comparison of relative modification rates in different genomic regions, or visualization of general modification profiles across regions of interest. In the extract operation, modification data is tabulated in a given genomic region for each read, reformatting single-read data for more efficient downstream processing. Both operation branches utilize modkit, an optimized tool created and published by ONT for parsing modified bases stored in BAM format ^28^. Outputs from these initial parsing operations are saved as compressed intermediate files to enable reuse in downstream analyses. The package provides methods for producing preset plots from these intermediate files, covering several common use cases for visualization of modified base data. Additionally, the methods for loading parsed data are made available to the user to enable the building of custom python analysis pipelines.

### Integrated multimodal analysis with *dimelo-toolkit*

We extended the original DiMeLo-seq protocol^13,14^ to make a multimodal version of the method, adapting it to enable simultaneous targeting of protein-DNA interactions and chromatin accessibility using distinct methyltransferases (Figure 2a). Combining antibody-targeted methylation with existing untargeted methylation techniques for measuring accessibility^15–17^ enables measurement of protein-DNA interaction, chromatin accessibility, and endogenous methylation on the same single-molecules in a single experiment. To mark protein-DNA interactions, we use a fusion of the nonspecific deoxyadenosine methyltransferase Hia5 and the antibody-binding protein A (pA-Hia5) to deposit 6mA near targets of interest. To mark accessible chromatin, we use the GpC methyltransferase M.CviPI to deposit 5mC at accessible GpC motifs. Methylation events representing each distinct epigenetic feature can be read in tandem using modification-sensitive long-read sequencing technologies such as Nanopore sequencing.

**Figure 2.**
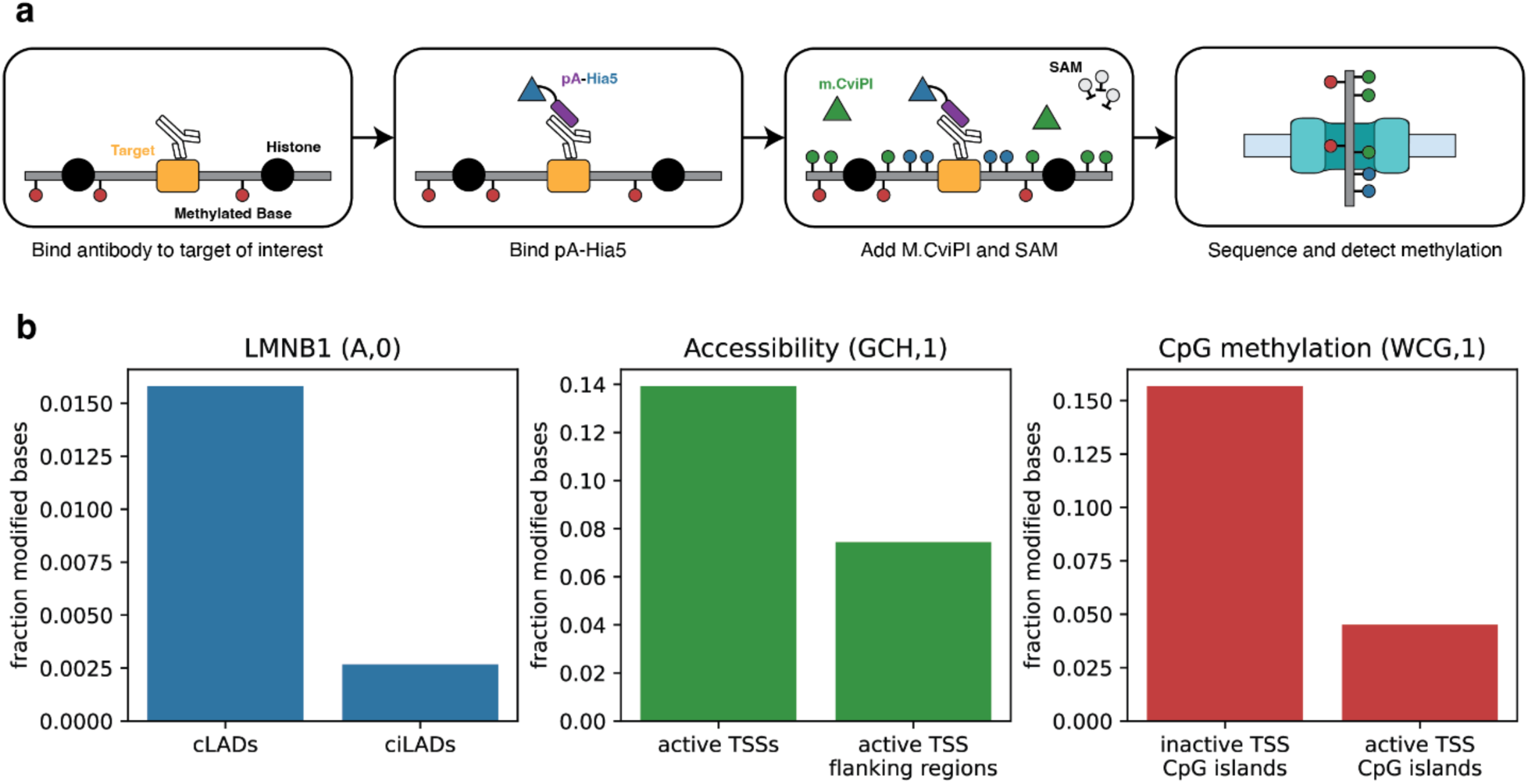
Multimodal DiMeLo-seq. **a**, Schematic of the multimodal DiMeLo-seq protocol for simultaneously mapping protein-DNA interactions, chromatin accessibility, and endogenous methylation. Epigenetic features are encoded on DNA using exogenous methylation, which can be read alongside endogenous methylation using long-read sequencing technologies. **b**, Barplots of modified base fractions in known on- and off-target regions for LMNB1 occupancy (blue), accessibility (green), and endogenous CpG methylation (red), measured simultaneously in an LMNB1-targeting multimodal DiMeLo-seq experiment in GM12878 cells. Pileups were performed with a modification probability threshold of 0.95.

To achieve optimal methylation performance, M.CviPI requires the reducing agent DTT, which is not present in the Hia5 buffer formulation described in the previously published DiMeLo-seq protocol^13,14^. It has been shown previously that antibodies can tolerate high concentrations of DTT and retain binding capability, and that Hia5 methylates efficiently in DTT-containing buffers^30^. We additionally found that Hia5 and M.CviPI can methylate simultaneously in Hia5 buffer supplemented with DTT, suggesting that there is minimal biochemical impact on enzyme performance when combining these methyltransferases (Supplementary Figure 1).

We applied the multimodal DiMeLo-seq method to simultaneously profile interaction sites of proteins of interest along with chromatin accessibility and endogenous CpG methylation in GM12878 cells. To validate the performance of this approach, we first targeted lamin B1 (LMNB1), which was previously used to optimize and validate the performance of DiMeLo-seq because lamina associated domains (LADs) are well-characterized genome-wide^6^. For each signal, we quantified the proportion of methylated motifs across all reads mapping to appropriate on-target and off-target regions and plotted the relative enrichment using the *plot_enrichment* method of *dimelo-toolkit* (Figure 2b). Relative methylation levels for each methylation signal followed the expected pattern of enrichment in on- and off-target regions, indicating that multimodal DiMeLo-seq can interrogate large-scale genome structure in the context of chromatin accessibility and endogenous methylation on single chromatin fibers.

To demonstrate the ability to visualize colocalization of protein interactions, accessibility, and endogenous CpG features, we chose to examine binding sites for CCCTC-Binding factor (CTCF). CTCF binding is inhibited by the presence of CpG methylation, and results in strong positioning of surrounding nucleosomes. We targeted CTCF with 6mA, measured chromatin accessibility with GpC methylation, and detected endogenous CpG methylation in GM12878 cells. We then investigated the resulting signals in aggregate at strong CTCF binding sites defined by ChIP-seq, using the *plot_enrichment_profile*, *plot_reads,* and *plot_read_browser* methods of dimelo-toolkit to visualize patterns in aggregate and on single molecules (Figure 3). We observed on-target increase of 6mA signal directly adjacent to the binding motif, inversely correlated to the CpG signal moving away from the binding sites (Figure 3a-b). All three signals exhibit a periodic pattern consistent with the periodic methylation of linker DNA between strongly-positioned nucleosomes. We additionally examined haplotype-specific patterns of CTCF binding, CpG methylation, and chromatin accessibility on single long reads. GM12878 has two X homologs, one of which has undergone X inactivation and remains inactive in all cells. We visualized CTCF binding patterns at the *Firre* locus, which is known to specifically bind CTCF on the inactive X chromosome^31,32^. With the multiplexed signals we can observe differential CTCF and accessibility patterns within the corresponding CpG landscape for both haplotypes. This analysis demonstrates the ability to detect and visualize three distinct epigenetic features with allele specificity on long, single molecules of genomic DNA.

**Figure 3.**
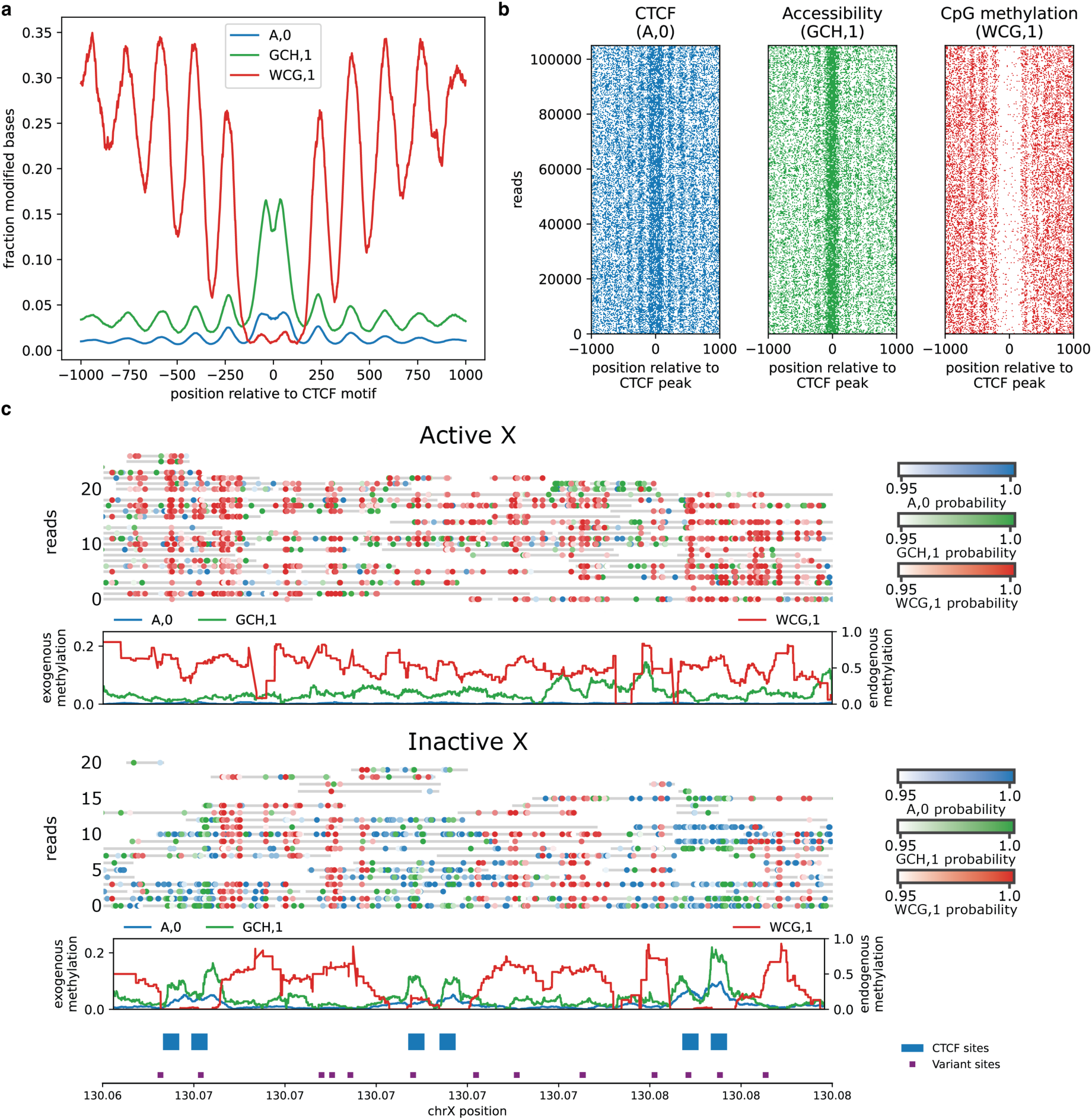
Multimodal analysis of CTCF binding sites. **a**, Enrichment profiles describing patterns of CTCF occupancy (blue), accessibility (green), and endogenous CpG methylation (red), as measured simultaneously in a CTCF-targeting multimodal DiMeLo-seq experiment in GM12878 cells. Modified bases reported with probability >0.95 were aggregated at 3,000 high-signal CTCF ChIP-seq peaks overlapping known CTCF motifs and plotted using a 25-bp rolling window. **b**, Single molecule plots for the reads used to generate the aggregate profiles in **(a)**. **c**, Single read and profile plots demonstrating methylation patterns for CTCF occupancy (blue), accessibility (green) and CpG (red) across active and inactive haplotypes at the Firre locus (chrX 130,064,000-130,080,000). Profile plots show fraction across all homolog reads smoothed with a 600bp rolling average. Annotations show variant sites used for read phasing and CTCF sites according to ChIP-seq peak calls.

We then expanded our analysis to demonstrate *dimelo-toolkit*’s support for other long-read assays and flexible data export for downstream analysis. We processed PacBio HiFi data from a Fiber-seq assay on the same GM12878 cell line from Vollger et al^33^ and exported all the different channels from both Fiber-seq and multimodal DiMeLo-seq in a genome-browser-compatible format using the *pileup_to_bigwig* method. This method allows the export of context-disambiguated modification profiles (e.g. WCG vs GCH), which is not natively supported in commonly used genome browsers such as IGV^34^. In Figure 4 we explore these tracks, along with data from short-read assays and ENCODE annotations, to show concordance and differences between the measurements and alignment to annotated features. This pipeline endpoint makes it easy to explore genomic loci of interest and cross-compare for more robust biological insights. Finally, we processed data from direct RNA sequencing of HEK293T cells from Hewel et al^35^ and used *dimelo-toolkit* to visualize patterns in N6-methyladenosine (m6A) and pseudouridine (ψ) across the transcriptome (Supplementary Figure 2). *dimelo-toolkit* enabled us to explore sequence context and functional enrichments that are expected based on the literature^36–39^ and visualize single-read modification patterns and colocalization to leverage the unique value of long-read sequencing.

**Figure 4.**
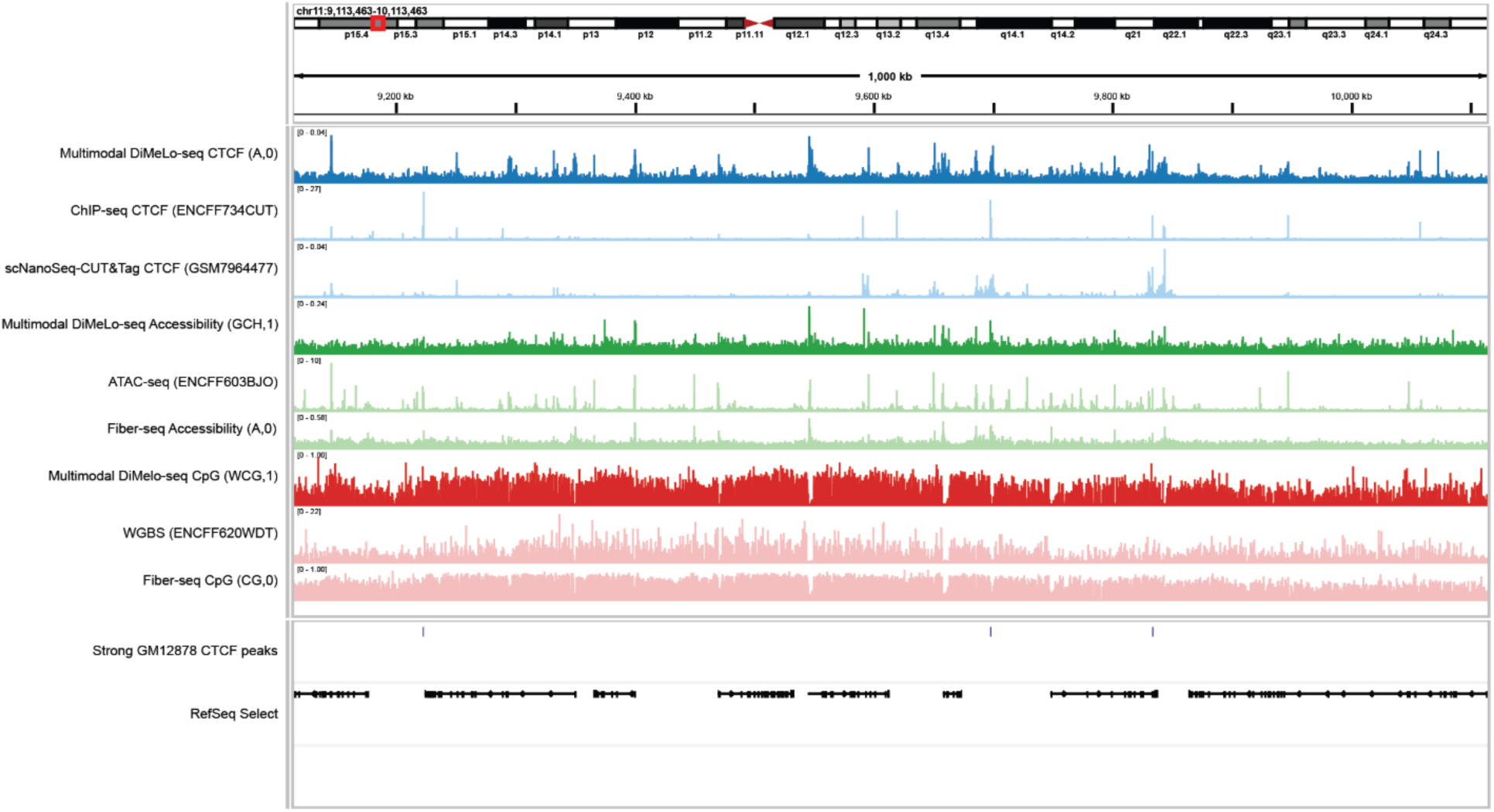
Genome browser comparison of multimodal DiMeLo-seq and orthogonal measurements. Aggregate browser traces from a CTCF-targeting multimodal DiMeLo-seq experiment in GM12878 cells (dark tracks) alongside reference data from comparable orthogonal measurements (ChIP-seq, scNanoSeq-CUT&Tag, ATAC-seq, WGBS, and Fiber-seq; light tracks), visualized in a representative 1,000 kb region of chromosome 11. Multimodal DiMeLo-seq pileups with a methylation probability threshold of 0.95 were exported in bigWig format using dimelo-toolkit.

### *dimelo-toolkit* facilitates assessment of bias in multimodal assays

Increasing the number of nucleotide modification signals creates unexpected technical problems which must be addressed to ensure accurate interpretation of the multimodal assays discussed here. First, if two or more signals use the same type of modification in different sequence motifs to represent distinct epigenetic features, care must be taken to exclude ambiguous motifs which could be interpreted as coming from either feature. For example, in the multimodal assay used in this study, methylation at GCG motifs can ambiguously be interpreted as both biologically relevant endogenous CpG methylation and exogenous GpC methylation representing chromatin accessibility; GCG motifs must therefore be omitted from analysis. Second, basecallers are usually not trained on data containing multiple types of exogenous modifications. Nanopore sequencing devices generate electrical traces representing multiple nucleotides passing through the pore at any given time. The presence of unexpected modifications in the pore may therefore confuse the basecaller, impacting modification detection in a model-specific way. For this assay, heavy GpC methylation is an out-of-distribution signal which empirically affects 6mA and CpG methylation levels reported by the models evaluated in this study.

To assess the impacts of adding a GpC methylation step to the existing DiMeLo-seq protocol, we first evaluated the effect of different M.CviPI incubation conditions on the reported signals for CpG and 6mA in a CTCF-targeting DiMeLo-seq experiment. We used the *plot_enrichment_profile* and *read_vectors_from_hdf5* functions to visualize all three signals as enrichment profiles and modification probability distributions and make direct comparisons between conditions (Figure 5, Supplementary Figure 3). With appropriate motif disambiguation, CpG signal was overall unperturbed by the addition of GpC methylation. 6mA probability distributions were notably perturbed in the presence of GpC methylation, reducing the separation between high- and low-confidence calls. However, targeted 6mA signal maintained clear peak structure and enrichment over baseline in enrichment profiles for all conditions. We selected a protocol which balances the quality of the targeted 6mA signal with enrichment for GpC accessibility and histone patterning, to maximize the useful biological insights that can be drawn from the data. To select appropriate disambiguated motifs for CpG and GpC signals for a multimodal application, we used the *plot_enrichment_profile* method to evaluate 5mC methylation detection in a variety of sequence contexts (Supplementary Figure 4). We found that the motifs with the best performance were sometimes stricter than those suggested by the methyltransferase specificities, supporting the notion that out-of-distribution modifications can perturb modification calling for nearby sites. These results led us to select a set of sequence motif specifications that minimized the overlap between accessible and endogenous methylation signals. Finally, we used the *read_vectors_from_hdf5* function to load python vectors for modification scores within reads localized to on- and off-target regions, allowing us to easily visualize the distributions of modification probabilities reported by the *dorado* basecaller (Supplementary Figures 5-6) and how these distributions shift in different contexts. This aided us in setting reasonable modification probability thresholds. By allowing for flexible data processing and visualization, dimelo-toolkit enabled us to differentiate biologically meaningful signal from technical bias for our multiplexed assay. The experimental conditions, motif specifications, and modification thresholds uncovered in this analysis were used throughout this study.

**Figure 5.**
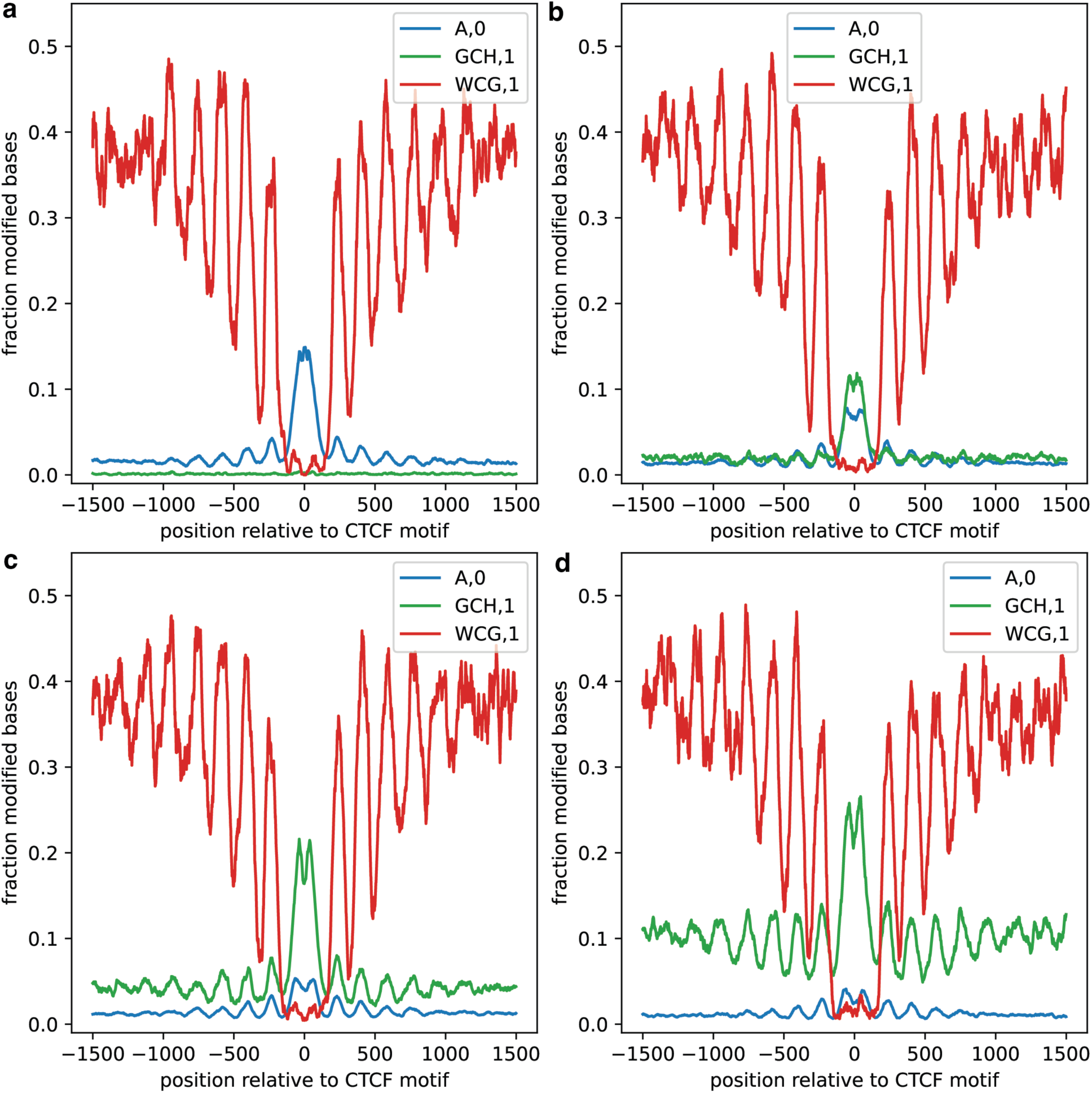
Evaluation of M.CviPI incubation protocols for multimodal DiMeLo-seq experiments. **a-d**, Enrichment profiles describing CTCF occupancy (blue), accessibility (green), and endogenous CpG methylation (red), as measured simultaneously in CTCF-targeting multimodal DiMeLo-seq experiments in GM12878 cells. M.CviPI incubation protocols varied across each of the four conditions: **(a)** no M.CviPI incubation, **(b)** incubation with 0.12 U/μl M.CviPI for the final 15 minutes of Hia5 incubation, **(c)** incubation with 0.12 U/μl M.CviPI for the full 2 hours of Hia5 incubation, and **(d)** incubation with 0.4 U/μl M.CviPI in a separate 15 minute incubation using M.CviPI buffer as adapted from the nanoNOMe-seq protocol^17^. **(c)** matches the protocol chosen for the multimodal DiMeLo-seq experiments in the remainder of this publication.

## Discussion

Here, we present *dimelo-toolkit*, a python package to facilitate analysis of assays reliant on long-read modification-aware nucleic acid sequencing technologies, such as Oxford Nanopore or PacBio HiFi. A key use case is to assist in developing and processing data from assays that involve adding exogenous nucleotide modifications to encode biological signals such as protein-DNA binding and chromatin accessibility. Our software aids users by providing a streamlined pipeline for downstream analyses and ready-made tools for producing both exploratory and publication-ready plots. We used this software package to analyze data from a new version of the DiMeLo-seq protocol which allows us to measure protein-DNA interactions and accessible chromatin on single DNA molecules in a single assay. We further demonstrated that *dimelo-toolkit* can be used to analyze and cross-compare data from many different protocols and chemistries, such as Fiber-seq sequenced with PacBio and Oxford Nanopore direct RNA sequencing.

When analyzing data from long-read modification assays, it is important to consider factors such as the selection of genomic regions of interest, read filtering strategies, selection of thresholds for modified basecall probabilities, and specification of modified sequence contexts. Providing an end-to-end python solution for this data makes it simple for researchers with varying levels of computational proficiency to build analysis pipelines and explore these parameters, helping users rapidly iterate, test hypotheses, and make informed data-driven assay development decisions. This process can be challenging for users because each factor cannot be considered independently. Score distributions, and thus appropriate thresholds, are impacted by sequence and modification contexts. Furthermore, protocol optimization results and disambiguation motif results can change based on the threshold used. *dimelo-toolkit* is therefore a key tool to use in this fundamentally iterative process.

Additionally, as new experimental methods are developed, a need arises for novel analytical methods for making meaningful biological inference. Providing direct access to the underlying data in python facilitates prototyping and development of tools to measure the distribution of data coming out of an assay and about the statistical confidence of biological conclusions. For example, exposing modified and unmodified counts from specified regions and sequence motifs makes it trivial to calculate p-values for differential states across conditions, paving the way for development of peak callers, comparing between cellular or chromosomal states, or integrating targeted and untargeted signals.

As new targets are explored with novel protocols for highly multiplexed analysis of epigenetic regulatory factors, *dimelo-toolkit* will accelerate data exploration and pipeline building through a unified framework for both one-click and bespoke analysis. By enabling researchers to better work with and understand multimodal long-read epigenetic data, this toolkit stands to accelerate the field and build towards a deeper understanding of regulatory genomics.

## Methods

### Methyltransferase functional validation assay

Assays were performed on 100 ng of a 360 bp dsDNA oligo containing a single restriction site each for AciI and MboI. Restriction sites were spaced such that unique combinations of methylation and digestion events result in fragments of unique lengths distinguishable using a High Sensitivity D1000 Tapestation kit. Methylation reactions using three different buffer conditions: 1X GC Reaction Buffer (NEB), DiMeLo-seq Activation Buffer (15 mM Tris, pH 8.0, 15 mM sodium chloride, 60 mM potassium chloride, 1 mM EDTA, pH 8.0, 0.5 mM EGTA, pH 8.0, 0.05 mM spermidine, 0.1% BSA), and DiMeLo-seq Activation Buffer containing 10mM DTT. All reactions were carried out in 30 μl volumes containing 800 µM SAM and incubated for 2 hours at 37 °C with SAM replenishment of 800 µM at the 1 hour mark. Following methyltransferase incubation, each condition was subjected to a restriction enzyme digestion for 2 hours in CutSmart buffer at 37 °C using AciI and MboI (NEB). A control sample of unmethylated oligo was subjected to the same digestion protocol. All digestion products were resolved using a High Sensitivity D1000 Tapestation kit.

### DiMeLo-seq + mGpC accessibility

Unfixed GM12878 cells were treated as described in past work, with the following modifications to accommodate the addition of an exogenous accessibility signal. Following primary and secondary antibody incubations, nuclei were resuspended in 100 µl of DiMeLo-seq Activation Buffer with the addition of 10mM DTT and 12 U of M.CviPI (NEB). Below, we provide the full protocol as adapted from previous publications^13,14^.

GM12878 cells (GM12878, Coriell Institute; mycoplasma tested) were maintained in RPMI-1640 with l-glutamine (Gibco, 11875093) supplemented with 15% FBS (VWR 89510-186) and 1% penicillin–streptomycin (Gibco, 15070063) at 37 °C in 5% CO_2_. After pelleting cells at 300g for 5 minutes and washing with PBS, nuclei were isolated by resuspending cells in 1 ml of Dig-Wash buffer (0.02% digitonin, 20 mM HEPES-potassium hydroxide buffer, pH 7.5, 150 mM sodium chloride, 0.5 mM spermidine, 1 Roche cOmplete EDTA-free tablet (11873580001) per 50 ml buffer and 0.1% BSA) and incubating on ice for 5 min. The resulting nuclei suspension was spun down at 4 °C at 500g for 3 min. All subsequent nuclei spins were performed with these same conditions, and all steps involving pipetting nuclei were performed with wide-bore tips. The supernatant was removed, and the pellet was gently resolved in 200 μl Tween-Wash (0.1% Tween-20, 20 mM HEPES-potassium hydroxide, pH 7.5, 150 mM sodium chloride, 0.5 mM spermidine, 1 Roche cOmplete EDTA-free tablet per 50 ml buffer and 0.1% BSA) containing the primary antibody at a 1:50 dilution. Antibodies targeted the following: LMNB1 (ab16048), CTCF (targeting C terminus, ab188408). Primary antibody incubation took place at 4 °C on a rotator overnight. Nuclei were then pelleted and washed twice with 0.95 ml Tween-Wash, with a 5 minute rotation at 4 °C between each spin. Following the second wash, the nuclei pellet was gently resolved in 200 μl Tween-Wash containing 200 nM pA–Hia5. Secondary antibody incubation took place at 4 °C on a rotator for 2 hours. Nuclei were then washed twice with 0.95 ml Tween-Wash in the same fashion as after the primary incubation. Nuclei were resuspended in 100 µl of DiMeLo-seq Activation Buffer containing DTT and M.CviPI (15 mM Tris, pH 8.0, 15 mM sodium chloride, 60 mM potassium chloride, 1 mM EDTA, pH 8.0, 0.5 mM EGTA, pH 8.0, 0.05 mM spermidine, 0.1% BSA, 10mM DTT, 12 U M.CviPI (NEB), and 800 μM SAM), followed by a 2 hour incubation at 37 °C with SAM replenishment of 800 µM at the 1 hour mark. DNA was extracted using the NEB Monarch Spin gDNA Extraction Kit (T3010S) and quantified using the Qubit 1x dsDNA BR Assay Kit (Q33265).

### Nanopore library preparation and sequencing

For each sample, 3 µg DNA was input into library preparation using one of the following kits: (1) Ligation Sequencing Kit V14 (ON SQK-LSK114) for single-library experiments, or (2) Native Barcoding Kit 24 V14 (ON SQK-NBD114.24) for optimization experiments and paired controls. For preparations using both kits, the protocol was performed as described in the gDNA version of the documentation, with the increased incubation times recommended in the original DiMeLo-seq protocol^13,14^. Sequencing was performed on an ONT P2Solo sequencer with v10.4.1 flow cells (ON FLO-PRO114M) with MinKNOW software (versions ranging from 24.06.14 to 24.11.10).

### Basecalling and postprocessing

Basecalling for DiMeLo-seq data was performed using ONT dorado^40^ (v1.0.2), using super-accuracy models for canonical bases, 5mC_5hmC, and 6mA (model complex: sup@v5.0.0,5mC_5hmC@v3,6mA@v3). Post-processing of basecalled data (trimming, aligning, demultiplexing, and summarization) was also performed using dorado, using base settings. Basecalled reads were mapped to the T2T-CHM13v2.0 reference sequence^41^. Uniquely-mapping reads of high average quality were selected based on the dorado summary (alignment_mapq == 60, mean_qscore_template > 10) and subsequently filtered using samtools^42^ to arrive at the final BAM files. For all analyses, bases with a methylation probability score >0.95 were assigned as methylated.

Haplotype phasing of DiMeLo-seq data was performed using WhatsHap (v2.6)^43^. Variant file for NA12878 was obtained from https://hgdownload.soe.ucsc.edu/gbdb/hg38/platinumGenomes/.

Basecalling and read processing for direct RNA data followed the setting described in the publication from which we sourced the data^35^, but upgrading to ONT dorado (v1.1.1). We chose the same super-accuracy model specification for canonical bases, m6A, and pseU (model setting commands: rna004_130bps_sup@v5.0.0 --modified-bases m6A pseU --estimate-poly-a). To uniquely map all of the data including rRNA tandem repeat transcripts and align in a splicing-aware fashion, basecalled reads were mapped to the hs1-rRNA v1.0 reference sequence^44^ with the dorado aligner, using --mm2-opts “-x splice:hq”.

### Genomic data file manipulation

Liftover operations from grch38 to T2T-CHM13v2.0 for publicly available files (BED, bigWig, etc.) were performed using the UCSC command line utilities^45^. Liftover operations from grch38 to T2T-CHM13v2.0 for publicly available VCF files were performed using picard (v 3.0.0)^46^. Additional operations (e.g. region merging, filtration, etc.) were performed using BEDtools^47^ (v2.31.1) and custom code available on Github.

### LMNB1 data analysis

Reference TSSs^48,49^ were ranked by expression levels using GM12878 bulk RNA-seq data (ENCFF978HIY, ENCODE Project Consortium). Top quartile TSSs by expression were labeled as active and bottom quartile TSS were labeled as inactive. Accessible target regions were defined as 1000 bp windows centered at active TSSs, and inaccessible target regions were defined as two 500 bp windows flanking each accessible target region. High and low CpG target regions were respectively defined as CpG islands^48^ located within 1000 bp of inactive and active TSSs. On- and off-target LMNB1 regions were respectively defined as cLADs and ciLADs identified by Altemose et al^6^.

### CTCF data analysis

Strong CTCF binding sites were defined as the top 3000 peaks from a set of reference GM12878 ChIP-seq peaks (ENCFF797SDL, ENCODE Project Consortium) which overlapped known CTCF motifs^50^. On-target CTCF sites were defined as 400 bp windows centered at strong CTCF binding sites, and off-target CTCF sites were defined as two 1300 bp windows flanking each on-target CTCF site.

### Direct RNA data analysis

Ribosomal RNA annotations were taken from George et al^44^. Stop codons were identified for transcripts genome wide using the NCBI RefSeq annotation for hs1, filtering for curated RefSeq transcripts (NM_/NR_ prefixes), and then identifying the 3’ end for the final CDS for each gene along the relevant strand. This yielded 66,619 sites genome-wide in a bed file with corresponding strand information.

## Data Availability

Raw .pod5 files and basecalled and aligned BAM files available upon request for GM12878 LMNB1 antibody+pA-hia5+M.CviPI (Figure 2), GM12878 CTCF antibody+M.CviPI high read coverage (Figure 3,4), and CTCF antibody+pA-hia5+M.CviPI incubation time experiments (Figure 5).

## Code Availability

*dimelo-toolkit*, along with installation instructions, documentation, and tutorials, can be found at https://github.com/streetslab/dimelo-toolkit. Code and instructions for generating the figures for this paper can be found at https://github.com/streetslab/multimodal-dimelo.

## Acknowledgments

Research reported in this publication was supported by the National Human Genome Research Institute and the National Institute of General Medical Sciences of the National Institutes of Health under award number R01HG012383 to A.S. O.D.L. was supported by the National Science and Engineering Research Council of Canada. A.S. and N.A. are Chan Zuckerberg Biohub – San Francisco Investigators. A.S. is supported by the Chan Zuckerberg Initiative SDL Award and the Harvey and Leslie Wagner Foundation. N.A. is an HHMI Hanna H. Gray Faculty Fellow and a Pew Biomedical Scholar. The authors would like to thank Karen Miga, Sofia Lundqvist, Denise Robles, Juliana Lee, and Pragya Sidhwani, Gary Karpen, and Arthur Rand for testing the software package and providing helpful discussion.

## Author Contributions

J.M., O.D.L., and A.S. designed the study. J.M. and A.M. performed the sequencing assays and experiments. J.M. and O.D.L. developed the software package, performed the analysis, generated figures, and wrote the manuscript. N.G. and M.R. beta tested the software package and helped conceive of new features. J.P.S. provided essential materials. A.F.S., N.A., N.M.I., F.D.U., and A.S. supported the work and provided guidance. A.S. supervised the study.

---

**Supplementary Figure 1.**
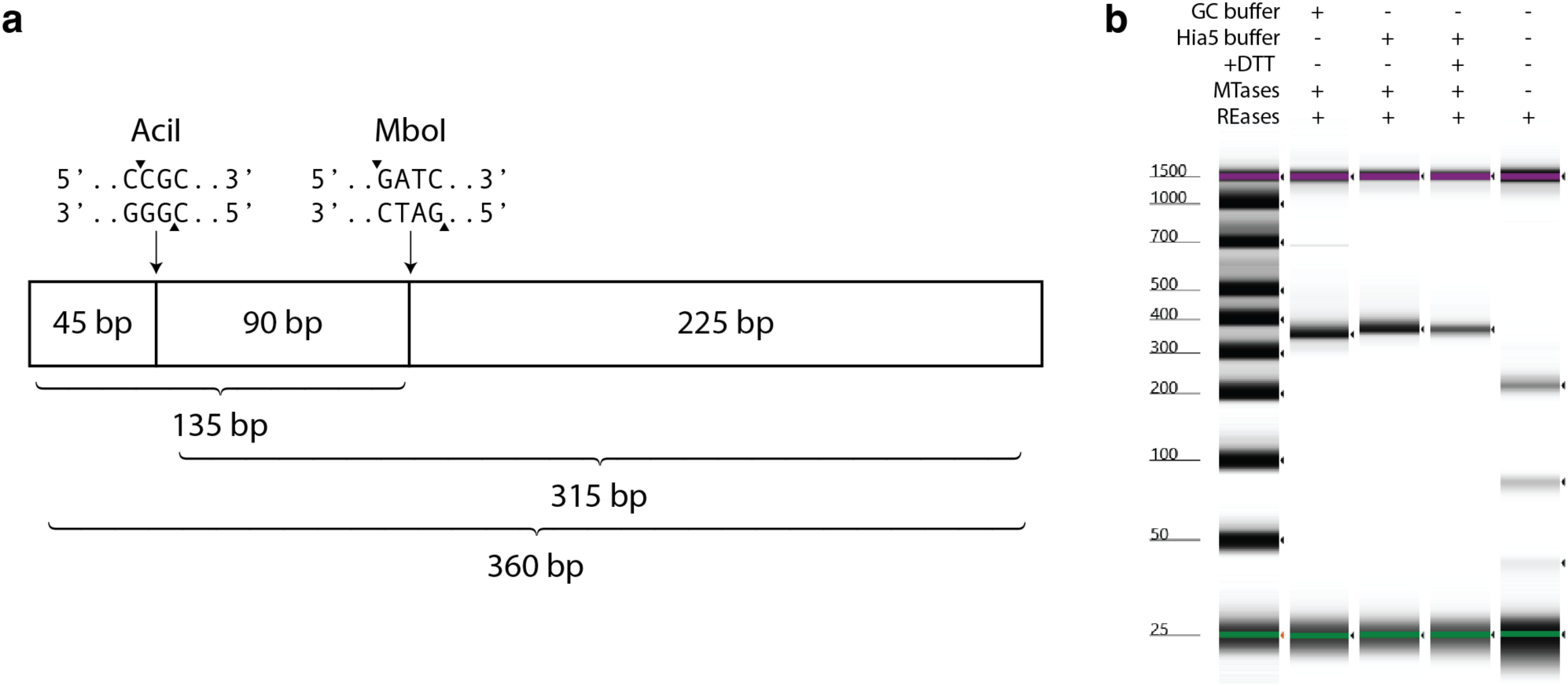
Validation of methyltransferase function using restriction enzyme digestion. **a,** Diagram of DNA template used for digestion assay. A 360 bp long unmethylated template was designed to contain a single restriction site each for AciI and MboI. AciI activity is blocked by cytosine methylation and MboI activity is blocked by adenine methylation. Restriction sites were placed such that unique combinations of methylation and digestion events result in fragments of unique lengths. **b,** Restriction enzyme digestion experiment testing methylation buffer conditions. DNA template from **(a)** was incubated with both M.CviPI and pA-Hia5 for 2 hours in different buffers, then subjected to restriction digest by AciI and MboI. GC buffer is NEB GC Reaction Buffer supplied with M.CviPI. Hia5 buffer is standard DiMeLo-seq methylation buffer. +DTT represents the addition of 10mM DTT. MTases and REases represent the addition of methyltransferases and restriction enzymes respectively.

**Supplementary Figure 2.**
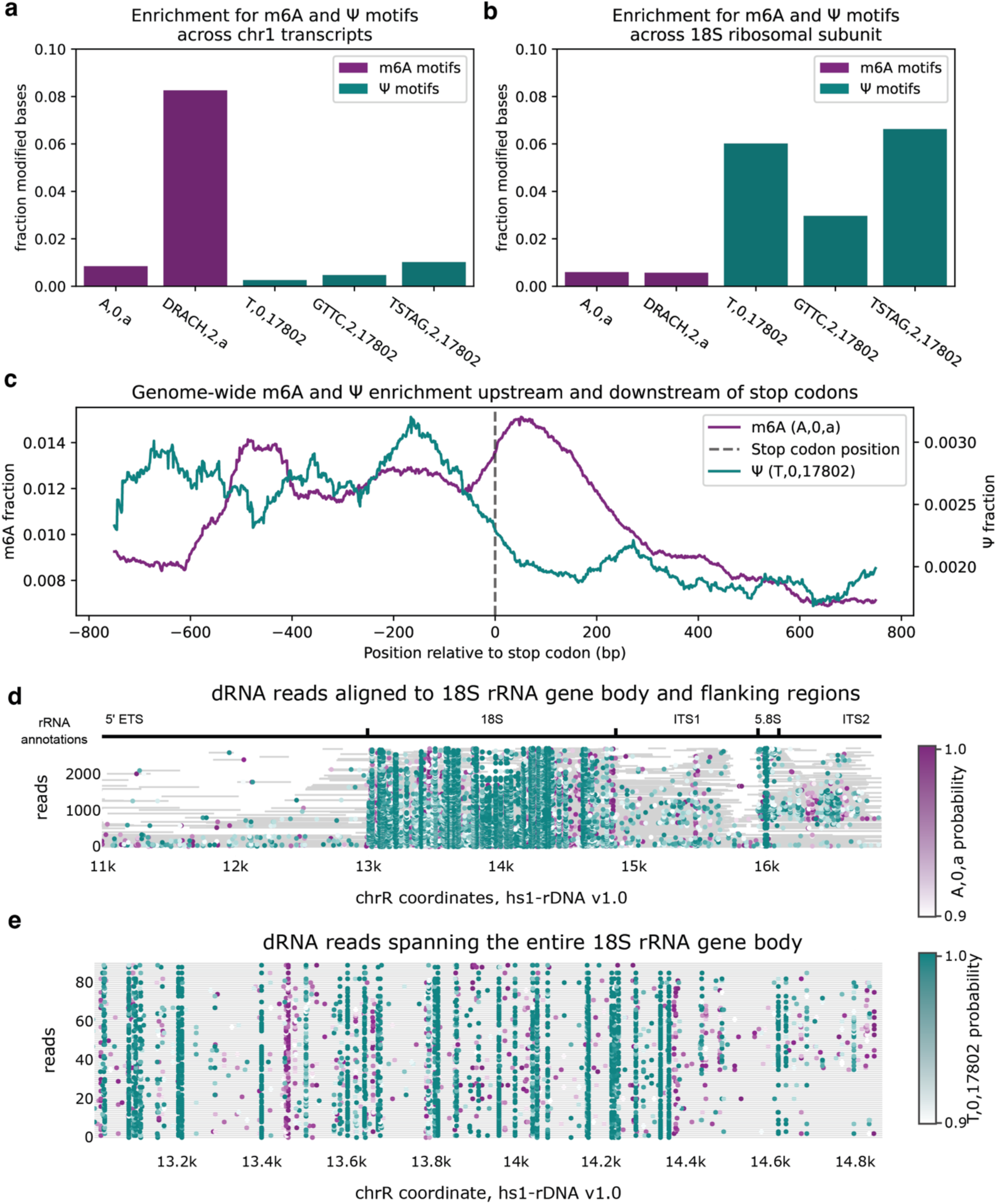
Demonstration of direct RNA modification analysis using dimelo-toolkit. **a**, Enrichment for RNA modifications for transcripts aligning to chr1. m6A and pseudouridine (ψ) are represented with modification codes ‘à and ‘17802’ respectively. chr1 transcripts exhibit the expected enrichment for m6A in DRACH motifs present on mRNA^1^, and expected enrichment for ψ in GUUC (GTTC) and TSTAG (USUAG) motifs^2^. **b**, Enrichment for RNA modifications for transcripts aligning to the 18S ribosomal subunit. Unlike in chr1, there is little to no additional enrichment for specific motifs in the 18S subunit, i.e. DRACH is not elevated above A alone and GTTC and TSTAG are not elevated above T alone. These observations are in line with known ribosomal RNA biology: high ψ levels are necessary for ribosome function and modifications occur by different mechanisms in rRNAs as compared to mRNA^3^. **c**, Enrichment profile of m6A and ψ across mRNAs, centered at the stop codon. m6A shows the expected enrichment peak immediately downstream of the stop codon in the 3’ UTR across 66,619 annotated CDS regions across the genome^4^. ψ conversely shows expected enrichment only in the CDS and not the 3’ UTR^2^. **d**, Single read browser showing aggregated reads across rRNA satellite regions. ψ positions are relatively consistent across the 18S rRNA. Many reads extend upstream into external transcribed spacer (ETS) and downstream into internal transcribed spacer (ITS1) and 5.8S rRNA (5.8S). This is in line with formation of the 47S precursor, as described in the literature^3^. **e**, Reads fully spanning the 18S rRNA (1864nt) allow us to see different single-read ψ and m6A colocalization patterns. Sorting is by read end coordinate, thus reads extending further into the internal transcribed spacer or potentially the 5.8S subunit show up higher on the plot. These reads exhibit different ψ and m6A near the end of the 18S subunit region, potentially indicating intermediate processing states.

**Supplementary Figure 3.**
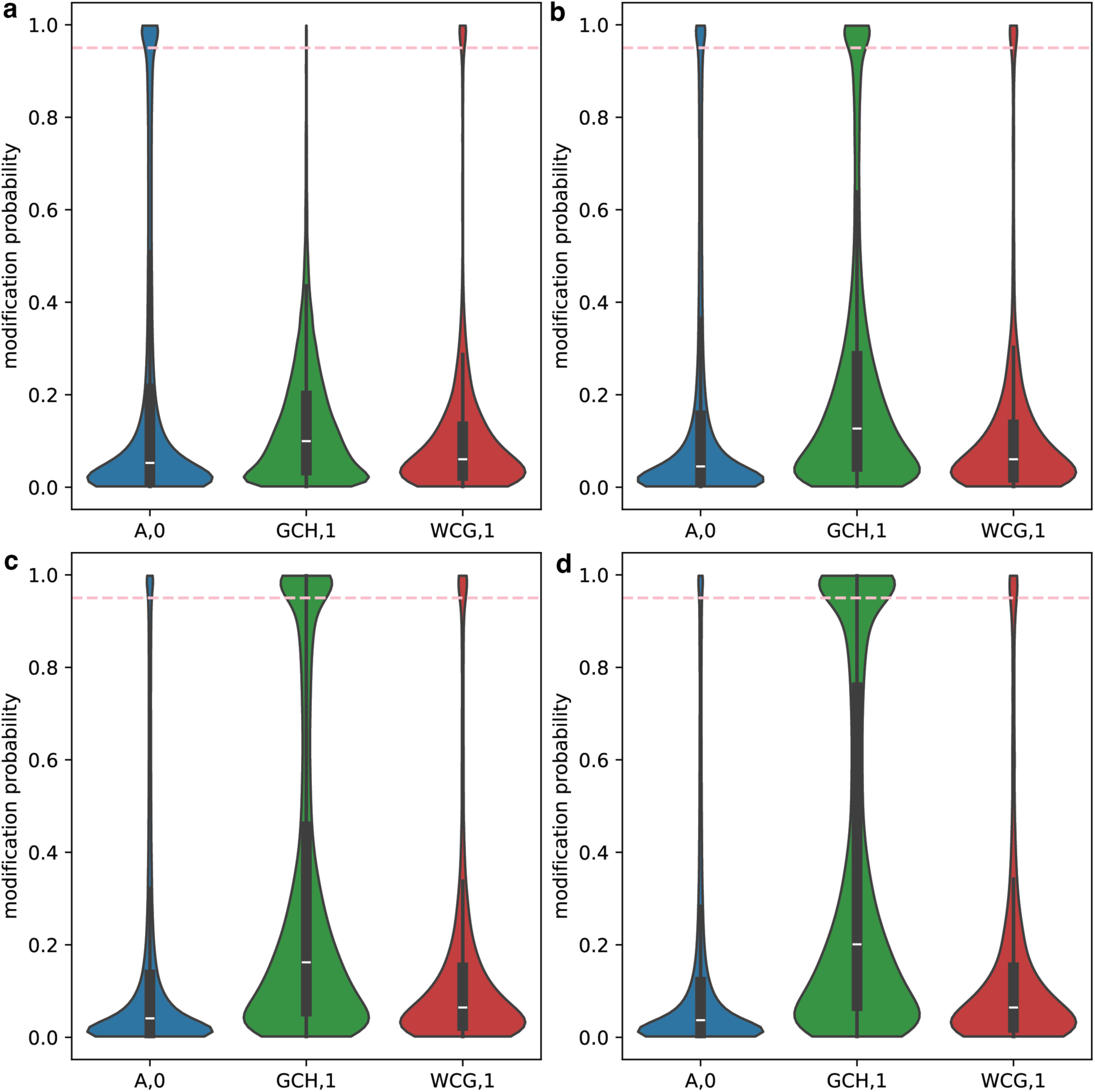
Effect of M.CviPI incubation protocols on multimodal DiMeLo-seq modification distributions. **a-d**, Distributions of modified base probabilities reported by the dorado basecaller for 6mA (blue; targeted to CTCF), 5mC in GC motifs (green; accessible chromatin), and 5mC in CG motifs (red; endogenous CpG methylation). The pink dashed line represents the modification probability threshold of 0.95 selected for subsequent analysis of this data. Distributions are calculated in 400 bp windows centered at 3,000 high-signal CTCF ChIP-seq peaks overlapping known CTCF motifs. M.CviPI incubation protocols varied across each of the four conditions: **(a)** no M.CviPI incubation, **(b)** incubation with 0.12 U/μl M.CviPI for the final 15 minutes of Hia5 incubation, **(c)** incubation with 0.12 U/μl M.CviPI for the full 2 hours of Hia5 incubation, and **(d)** incubation with 0.4 U/μl M.CviPI in a separate 15 minute incubation using M.CviPI buffer as adapted from the nanoNOMe-seq protocol^5^. **(c)** matches the protocol chosen for the multimodal DiMeLo-seq experiments in the remainder of this publication.

**Supplementary Figure 4.**
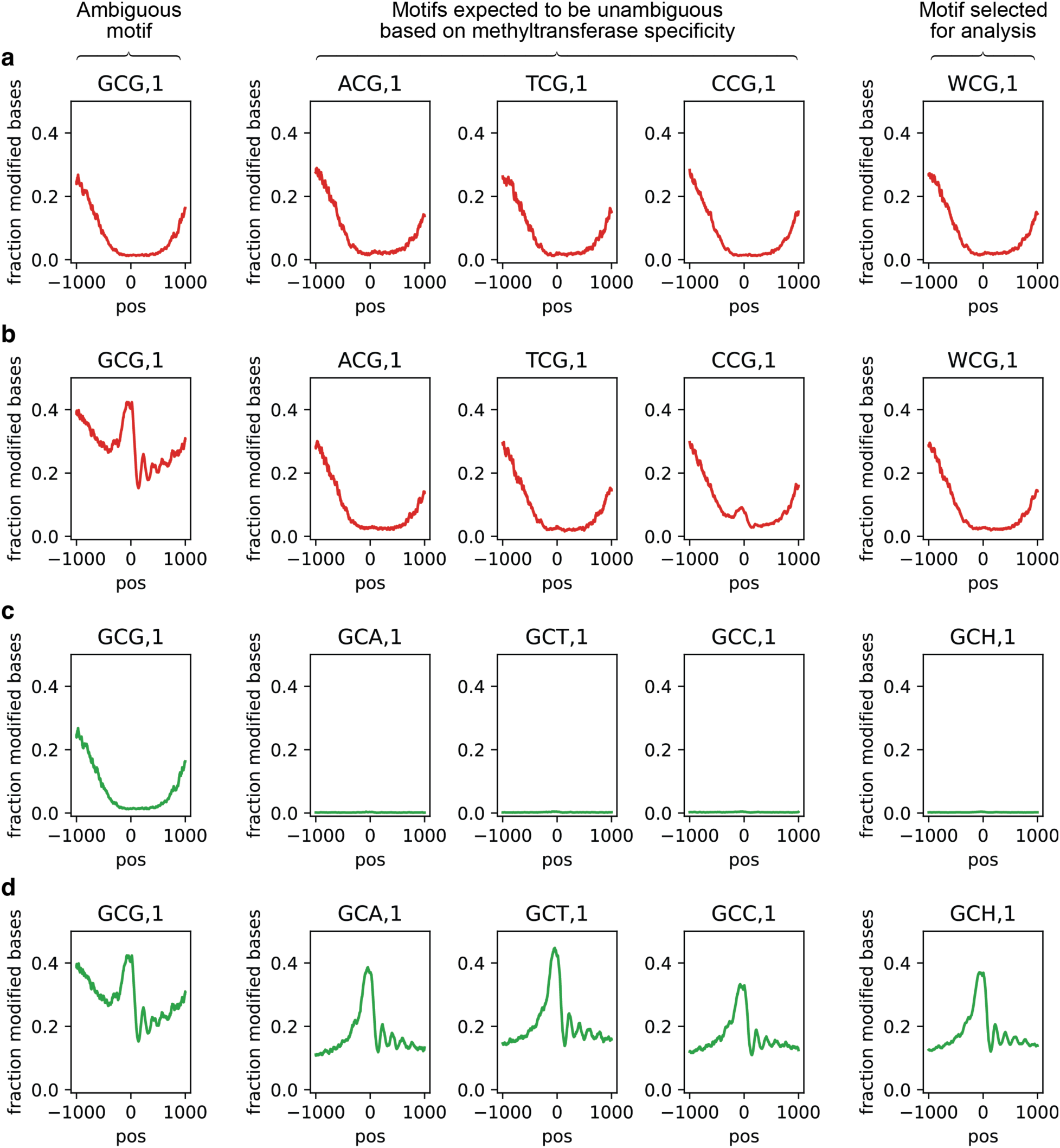
Motif disambiguation for 5mC detection in the presence of exogenous GpC methylation. **a-b**, CpG methylation profiles (red) for high-expression TSSs, at CG motifs with different preceding bases, in untreated **(a)** and M.CviPI-treated **(b)** conditions. **c-d**, GpC methylation profiles (green) for high-expression TSSs, at GC motifs with different succeeding bases, in untreated **(c)** and M.CviPI-treated **(d)** conditions. All profiles were generated using a modification probability threshold of 0.75 and a 25 bp rolling window. Subplot titles represent the motif codes used for each pileup operation. Left column represents the GCG motif which is shared between CG and GC methyltransferases. Middle columns represent motifs which are not expected to be shared between the methyltransferases. Right column represents the disambiguated motif selected for the analyses in this study.

**Supplementary Figure 5.**
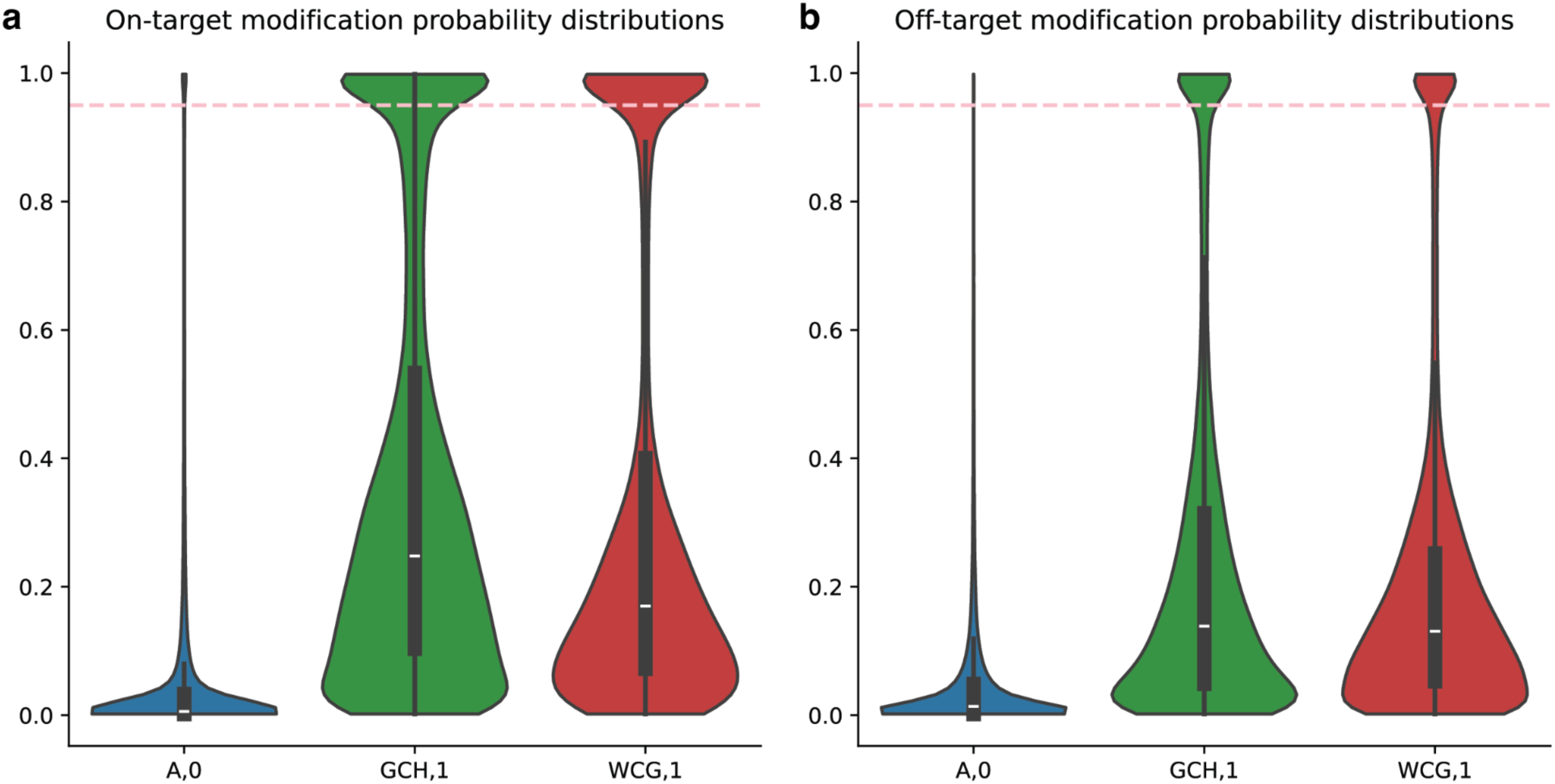
Modification probability distributions in LMNB1-targeting multimodal DiMeLo-seq experiment in GM12878 cells. **a-b,** Distributions of modified base probabilities reported by the dorado basecaller for 6mA (blue; targeted to LMNB1), 5mC in GC motifs (green; accessible chromatin), and 5mC in CG motifs (red; endogenous CpG methylation). The pink dashed line represents the modification probability threshold of 0.95 selected for subsequent analysis of this data. Distributions are shown for on-target regions **(a)** and off-target regions **(b)** for each methylation signal. On- and off-target regions are defined the same way as in Figure 2: for 6mA, on-target regions are cLADs and off-target regions are ciLADs. For 5mC in GC motifs, on-target regions are active TSSs and off-target regions are the flanking regions of those TSSs. For 5mC in CG motifs, on-target regions are CpG islands overlapping inactive TSSs and off-target regions are CpG islands overlapping active TSSs.

**Supplementary Figure 6.**
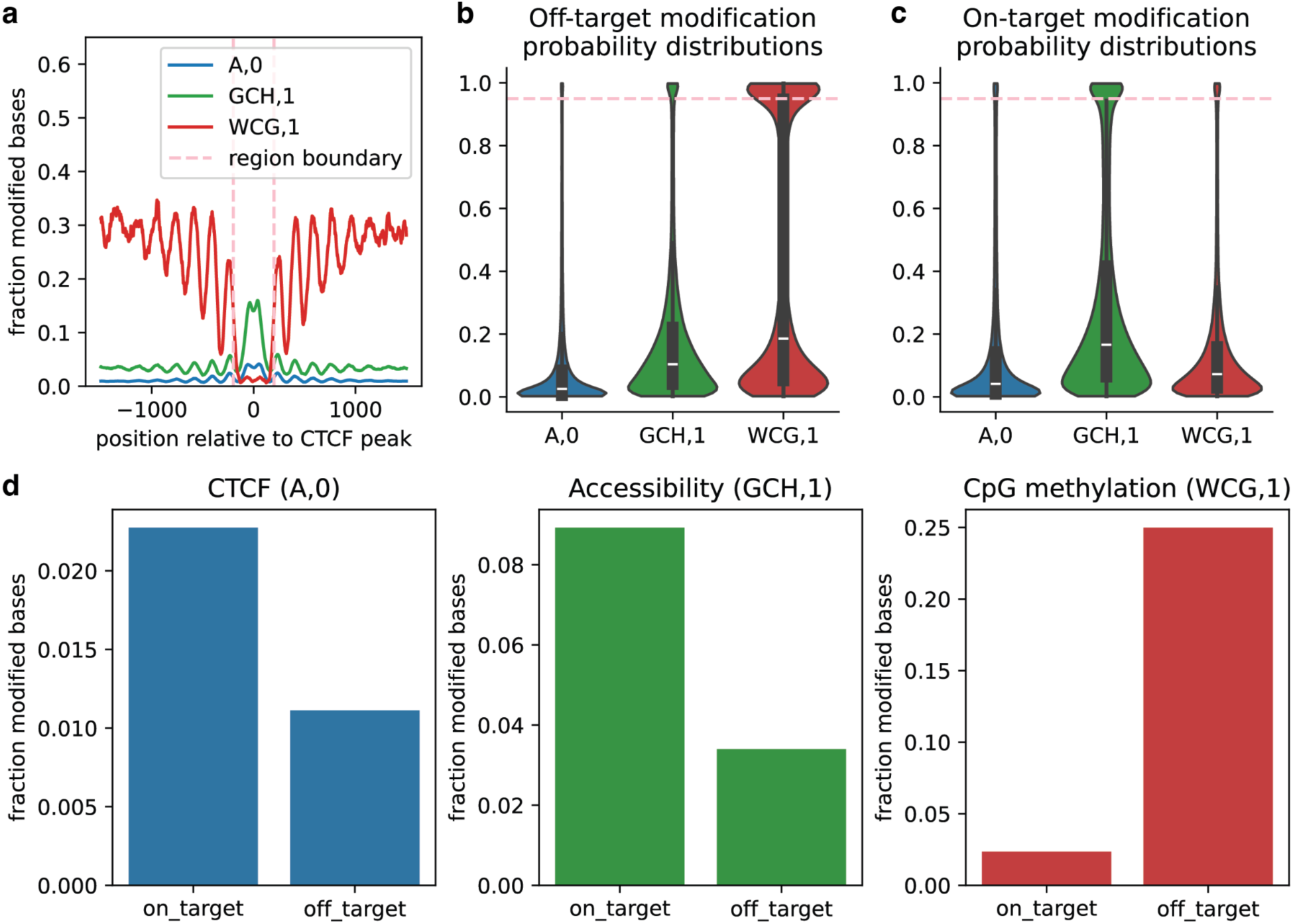
Modification probability distributions in CTCF-targeting multimodal DiMeLo-seq experiment in GM12878 cells. **a,** Enrichment profiles describing CTCF occupancy (blue), accessibility (green), and endogenous CpG methylation (red). Pink dashed lines represent the boundary defining on-target regions (+/- 200 bp from peak center) and off-target regions (1,300 bp flanking on-target regions on both sides). **b-c,** Distributions of modified base probabilities reported by the dorado basecaller for 6mA (blue; targeted to CTCF), 5mC in GC motifs (green; accessible chromatin), and 5mC in CG motifs (red; endogenous CpG methylation). The pink dashed line represents the modification probability threshold of 0.95 selected for subsequent analysis of this data. Distributions are shown in the on-target **(b)** and off-target **(c)** regions defined in **(a). d**, Barplots of modified base fractions in regions defined in **(a)** for modifications reported with probability >0.95.

